# Controlling CRISPR-Cas9 with ligand-activated and ligand-deactivated sgRNAs

**DOI:** 10.1101/323105

**Authors:** Kale Kundert, James E. Lucas, Kyle E. Watters, Christof Fellmann, Andrew H. Ng, Benjamin M. Heineike, Christina M. Fitzsimmons, Benjamin L. Oakes, David F. Savage, Hana El-Samad, Jennifer A. Doudna, Tanja Kortemme

**Affiliations:** Graduate Group in Biophysics, University of California San Francisco, San Francisco, CA, USA; UC Berkeley – UCSF Graduate Program in Bioengineering, University of California San Francisco, San Francisco, CA, USA; Department of Molecular and Cell Biology, University of California Berkeley, Berkeley, CA, USA; Bioinformatics Graduate Program, University of California San Francisco, San Francisco, CA, USA; Chemistry and Chemical Biology Graduate Program, University of California San Francisco, San Francisco, CA, USA; Department of Biochemistry and Biophysics, University of California San Francisco, San Francisco, CA, USA; Chan Zuckerberg Biohub, San Francisco, CA, USA; Howard Hughes Medical Institute, University of California Berkeley, Berkeley, CA, USA; Department of Bioengineering and Therapeutic Sciences, University of California San Francisco, San Francisco, CA, USA

## Abstract

The CRISPR-Cas9 system provides the ability to edit, repress, activate, or mark any gene (or DNA element) by pairing of a programmable single guide RNA (sgRNA) with a complementary sequence on the DNA target. Here we present a new method for small-molecule control of CRISPR-Cas9 function through insertion of RNA aptamers into the sgRNA. We show that CRISPR-Cas9-based gene repression (CRISPRi) can be either activated or deactivated in a dose-dependent fashion over a >10-fold dynamic range in response to two different small-molecule ligands. Since our system acts directly on each target-specific sgRNA, it enables new applications that require differential and opposing temporal control of multiple genes.

## Introduction

CRISPR-Cas9 has emerged as an immensely powerful system for engineering and studying biology due to its ability to target virtually any DNA sequence *via* complementary base pairing with a programmable single-guide RNA (sgRNA)[1]. This ability has been harnessed to edit genomes, repress[2] or activate[3, 4] gene expression, image DNA loci[5], generate targeted mutational diversity[6] and to modify epigenetic markers[7].

In addition to engineering CRISPR-Cas9 for diverse applications, there has also been broad interest in developing strategies to regulate CRISPR-Cas9 activity[8, 9]. Such strategies promise to mitigate off-target effects and allow the study of complex biological perturbations that require temporal or spatial resolution[9]. To date, most of the progress in this area has been focused on switching the activity of the Cas9 protein using chemical[10-16] or optical[17-19] inputs. A general issue with these approaches is that all target genes are regulated in the same manner, although this limitation can be addressed with orthogonal CRISPR-Cas9 systems[16, 20, 21].

An alternative but less explored strategy is to regulate the sgRNA instead of the Cas9 protein. Since the sgRNA is specific for each target sequence, controlling the sgRNA directly has the potential to independently regulate each target. This strategy has been approached using sgRNAs that sequester the 20 nucleotide target sequence (the spacer) only in the absence of an RNA-binding ligand[22, 23], ligand-dependent ribozymes that cause irreversible RNA cleavage[23, 24], ligand-dependent protein regulators recruited to the sgRNA to alter CRISPR function[12], and engineered antisense RNA to sequester and inactivate the sgRNA[25].

Here we describe a new method to engineer ligand-responsive sgRNAs by using RNA aptamers to directly affect functional interactions between the sgRNA, Cas9, and the DNA target (**Figure 1a**). In contrast to prior sgRNA-based methods, our approach can be used to both activate and deactivate CRISPR-Cas9 function in response to a small molecule. In addition, our approach requires only Cas9 and the designed sgRNAs. We further show that control of CRISPR-Cas9 function with our method is dose-dependent over a wide range of ligand concentrations and can be used to simultaneously execute different temporal programs for multiple genes within a single cell. We envision that this method will be broadly useful for regulating essentially all applications of CRISPR-Cas9-mediated biological engineering.

## Results and Discussion

We sought to insert an aptamer into the sgRNA such that ligand binding to the aptamer would either activate or deactivate CRISPR-Cas9 function. We envisioned that ligand binding could either stabilize or destabilize a functional sgRNA conformation — bound to Cas9 and the DNA target — over other competing states in the ensemble (**Figure 1a**). We chose the theophylline aptamer[26] as a starting point because it is well-characterized and has high affinity for its ligand, which is cell permeable and is not produced endogenously.

We first asked which sites in the sgRNA were most responsive to the insertion of the theophylline aptamer and which strategies for linking the aptamer to the sgRNA were most effective. We designed aptamer insertions at each of the sgRNA stem loops at sites that are solvent-exposed in the Cas9/sgRNA/DNA ternary complex[27] and exhibit various levels of tolerance to mutation[28]. These insertion sites are denoted the upper stem, nexus, and hairpin (**Figure 1b**). We tested three linking strategies aimed at stabilizing a functional sgRNA conformation in the presence of the ligand: (i) replacing parts of each stem with the aptamer, (ii) splitting the sgRNA in half and using the aptamer to bring the halves together, and (iii) designing strand displacements (i.e. sequences that allow for alternative base pairing in the *apo* and *holo* states) (**Figure 1c**, **Table S1**).

To test the resulting 86 designed sgRNAs, we used an *in vitro* assay to measure differential Cas9-mediated DNA cleavage in the presence and absence of theophylline. We identified theophylline-responsive sgRNAs for all three insertion sites (**Figure 1d**), with the most successful designs derived from the strand displacement linking strategy (**Table S1**). We confirmed that the activity of our designs depended on the concentration of theophylline, as would be expected if the ligand affects function through binding the aptamer-containing designed sgRNA (**Figure 1e**). In total, 10 designs were responsive to theophylline *in vitro*. For nine of these responsive designs theophylline addition activated CRISPR-Cas9 function, while for one design (#61) theophylline unexpectedly deactivated function. Interestingly, all of the theophylline-activated designs had the aptamer inserted into either the upper stem or the hairpin, while the theophylline-deactivated design had the aptamer inserted into the nexus (**Figure 1b,d**, **Table S1**). These findings suggested the exciting possibility of regulating CRISPR-Cas9 function with both ligand-activated and ligand-deactivated sgRNAs, depending on the aptamer insertion site.

We next sought to find designed sgRNAs that would function robustly in *E. coli.* To screen designed libraries using fluorescence-activated cell sorting (FACS), we changed to a cellular assay based on CRISPR-Cas9-mediated repression (CRISPRi) of super-folder green fluorescent protein (sfGFP) and monomeric red fluorescent protein (mRFP) (**Figure 2a**)[14]. The strongest rational designs exhibited only weak activity in the CRISPRi assay (**Figure S1**). Since the sequences linking the aptamer to the remainder of the sgRNA affected activity in our *in vitro* experiments, we designed libraries with randomized linkers of 4-12 nucleotides at all three insertion sites to broadly sample different aptamer contexts (**Figure 2b**, **Table S2**). To identify ligand-activated sgRNAs, we screened each library first for CRISPRi activity in the presence of theophylline, then second for lack of activity in the absence of theophylline. We then repeated that selection/counter-selection with a different spacer to avoid selecting sgRNA scaffold sequences that would be specific for a particular spacer (**Figure 2c**). To identify ligand-inhibited sgRNAs, we used an analogous four-step selection/counter-selection protocol that began by screening for activity in the absence of ligand. We validated the activity of the selected hits with a third spacer that was not used in any of the screens (**Table S3**). The most robust ligand-activated sgRNA variant (termed ligRNA^+^; i.e. sgRNA that is active in the + ligand state) and ligand-inactivated sgRNA (termed ligRNA^−^) showed 11x and 13x dynamic ranges that spanned 55% and 59% of the range achieved by the controls, with negligible overlap between the active and inactive populations (**Figure 2d,e**, **Table S4**).

The ligRNA^+^ construct derived from inserting the aptamer into the hairpin while randomizing the remainder of the hairpin, the 5 unpaired nucleotides at the apex of the nexus, and the region between the hairpin and the nexus. The hairpin became more GC-rich, but the base-pairing was conserved with the exception of a single mismatch. The apex of the nexus became complementary to the 5’ region of the aptamer. The region between the hairpin and the nexus remained AU-rich and unpaired (**Figure S2a**). Secondary structure predictions of ligRNA^+^ using ViennaRNA[29] (**Figure S2b**) are consistent with our intended mechanism, where ligand binding to the aptamer leads to strand displacements stabilizing the active sgRNA conformation.

The ligRNA^−^ construct derived from inserting the aptamer into the nexus while randomizing the nexus stem. The nexus stem was extended from 2 to 5 bp, but remained base-paired and GC-rich (**Figure S2a**). The mechanism underlying the ability of ligRNA^−^ to deactivate CRISPR-Cas9 function upon the addition of theophylline was unclear. ViennaRNA predictions of the lowest energy conformation for ligRNA^−^ were uninformative on the mechanism of ligand control, as they suggested that ligRNA^−^ adopts the same secondary structure in the presence and absence of theophylline (**Figure S2c**). However, we noticed that in the stem sequence selected in our screen, U95 in ligRNA^−^ (U59 in the crystal structure of the Cas9 ternary complex with DNA and RNA[27]) was conserved in 17 of the 20 isolated sequences (**Table S3**). This uracil makes specific hydrogen-bonding interactions with asparagine 77 in the ternary complex. In the sgRNA scaffold this uracil is always unpaired, but in ligRNA^−^ it is predicted to engage in a wobble base pair with G65 in the stem leading up to the aptamer (**Figure S2c**). These observations led us to hypothesize that ligand binding to the aptamer controls the extent to which U95 is unpaired, which in turn determines whether or not ligRNA^−^ interacts functionally with Cas9 and the target DNA. To test this hypothesis, we first designed strand-swapping mutations in the stem leading up to the aptamer (**Figure S3**). As expected, swapping U95 rendered ligRNA^−^ completely inactive, while swapping base pairs at the positions between U95 and the aptamer had only a mild effect (**Figure S3d**). We then modulated the strength of the base pairs between U95 and the aptamer. Consistent with the hypothesis that ligand binding to the aptamer decreases access to U95, we found that weaker base pairs were more repressing while stronger base pairs were more activating (**Figure S3e**). These results provide a possible explanation for how ligRNA^−^ deactivates CRISPR-Cas9 function in the presence of the ligand and suggest additional ways for ligRNAs to be tuned for specific applications.

Next, we tested whether the ligRNAs responded to increasing concentrations of theophylline in a dose-dependent manner in the cellular CRISPRi assay. We observed that the activities of both ligRNA^+^ and ligRNA^−^ were smoothly titratable and exhibited a nearly linear response over a large range of ligand concentration (**Figure 2f**). We note that the apparent EC_50_s of ligRNA^+^ and ligRNA^−^ (134.3±11.3 μM and 177.6±17.3 μM) are much higher than the *K*_*D*_ of the theophylline aptamer alone (320 nM)[30]. This discrepancy is common for RNA devices[31] and could be explained by the altered structural context of the aptamer embedded in an sgRNA sequence. Nevertheless, the linear concentration dependence of the ligRNAs demonstrates their utility for not only turning genes on or off, but also for precisely tuning their levels of expression.

Because RNA devices are known to be sensitive to sequence context[32], we tested ligRNA^+^ and ligRNA^−^ with 24 different spacers using the *in vitro* DNA cleavage assay (**Figure 2g**, **Figure S4**, **Table S5**). We found that both ligRNA^+^ and ligRNA^−^ respond to theophylline for the majority of the tested spacers (15 and 21 out of 24 spacers for ligRNA^+^ and ligRNA^−^, respectively). For the few spacers that did not function, we hypothesized that base-pairing of the spacer sequence with the aptamer might explain the lack of sensitivity to theophylline. To address this question, we predicted the affinity between each spacer and the aptamer (with its associated linker) for both ligRNAs using ViennaRNA[29] (**Figure S5**). For ligRNA^+^ constructs the correlation between the duplex free energy prediction and theophylline sensitivity was negligible. However, for ligRNA^−^ constructs increased predicted affinity of the spacer for the aptamer sequence correlated with a smaller change in Cas9-mediated DNA cleavage in response to theophylline. This analysis suggested that spacers with predicted affinity for the aptamer could interfere with switching of the ligRNA^−^ function, providing a useful design criterion for functional spacers. Taken together, these results suggest that ligRNAs should be capable of regulating most genes, especially those that can be targeted by multiple spacers.

To test how decreased expression levels would affect the ligRNAs, we replaced the strong constitutive promoter driving sgRNA expression (J23119) with a weak constitutive promoter (J23150) and repeated the CRISPRi assay. Both ligRNAs remained functional with the weak promoter, albeit with a somewhat narrower dynamic ranges (from a 10.2±0.7-fold to a 5.9±0.5-fold change upon theophylline addition for ligRNA^+^ and from a 16.2±1.0-fold to a 11.6±0.9-fold change for ligRNA^−^, **Figure S6**). Notably, the weak promoter shifted the dynamic ranges of both ligRNAs in the direction of increased gene expression, to the point where nearly full gene activation was achieved in the non-repressing state. These results suggest that the ligRNAs are able to repress at low expression levels, and that tuning promoter strength is useful for applications that require full gene activation (alternatively, a collection of ligRNA variants that shift the dynamic range is shown in **Figure S7**).

A key advantage of regulating CRISPR-Cas9 using the sgRNA instead of the protein is the ability to independently control different genes with different ligands in the same Cas9 system. To test this idea, we replaced the theophylline (theo) aptamer in the ligRNAs with the 3-methylxanthine (3mx) aptamer (the resulting sgRNA constructs were termed ligRNA^(+/−)3mx^). While the theophylline aptamer is recognized by both ligands, the 3-methylxanthine aptamer is specific to its ligand[33]. Since the aptamers differ in only one position, the replacement of the theophylline aptamer with the 3-methylxanthine aptamer was straightforward and led to a 3-methylxanthine-sensitive ligRNA^−^ variant without further optimization. (ligRNA^+^ also remained functional with the 3-methylxanthine aptamer but exhibited an undesirable albeit small ~2-fold response to theophylline, **Figure S8**). We used the two ligRNA^−^ variants to construct a system that expresses GFP upon addition of theophylline and expresses GFP and RFP when both ligands are added (Figure 3a). We then performed a timecourse where we sequentially activated, deactivated, and reactivated both reporter genes using ligRNAs and observed the expected temporal expression program (Figure 3b).

Taken together, ligRNAs provide control of both gene repression and gene activation and can be multiplexed for differential control of genes in the same system (Figure 3). While ligRNAs function robustly in bacteria, transferring them to eukaryotic systems will require further optimization (**Figure S9**, **Figure S10**) which could be achieved using methods similar to our optimization (Figure 2) of the initial rational designs (Figure 1). Nevertheless, there are already many useful applications for ligRNAs in bacteria. For example, many species of bacteria do not have facile genetic controls available, and ligRNAs provide such controls with a minimal footprint. Moreover, temporally controlled gene expression programs are thought to be important for key biological processes in bacteria[34], and ligRNAs provide a way to conduct large-scale screens to probe these programs and their role in the interactions between bacteria and their environments[35].

The study of subtle effects in complex biological systems will increasingly require the ability not just to probe individual genes, or to knock down different sets of genes, but to tune the expression of many different genes with fine temporal precision. ligRNAs provide this capability by adding ligand- and dose-dependent control of individual sgRNAs to the already powerful CRISPR-Cas9 technology.

**Figure 1:**
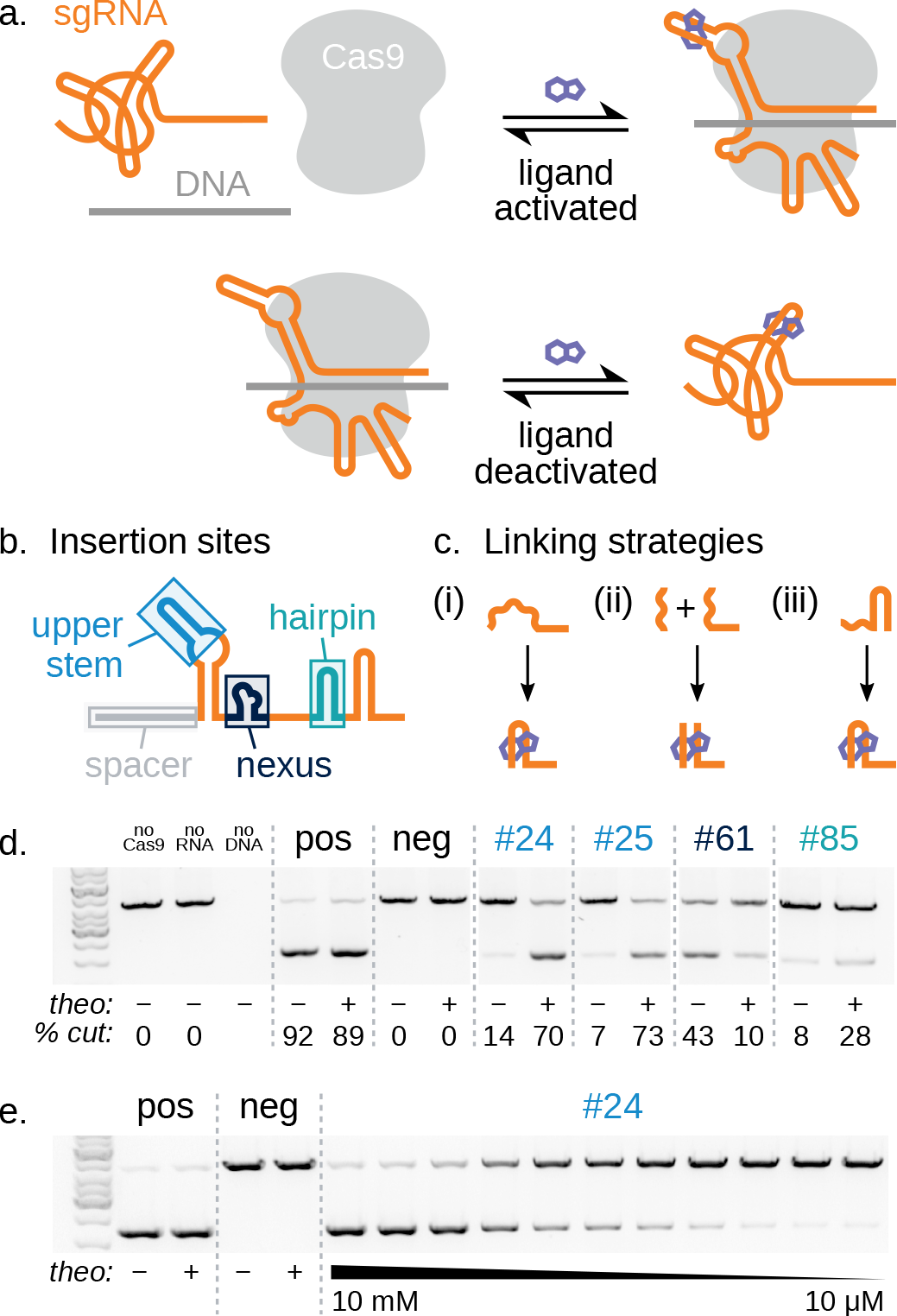
Design of ligand-controlled sgRNAs by inserting small molecule aptamers into the sgRNA. (**a**) Illustration of the design goal, where functional Cas9/sgRNA/target DNA complexes are stabilized either in the presence (**top**) or absence (**bottom**) of a small molecule ligand. (**b**) Aptamer insertion sites; sgRNA domains defined as in ref.[28]. (**c**) Strategies for linking the aptamer to the sgRNA: (i) stem replacement; (ii) induced dimerization; (iii) strand displacement (**Table S1**). (**d**) Efficiency of *in vitro* Cas9 cleavage of DNA in the presence and absence of 10mM theophylline for controls and selected designs. Design numbers refer to **Table S1** and are color-coded by aptamer insertion sites defined in (**b**). Percent cut values (bottom) are the average of at least two experiments. All data shown are from a single gel (some lanes are excluded for clarity). (**e**) Dose-dependence of cleavage efficiency in response to increasing concentrations of theophylline for a representative design (#24).

**Figure 2:**
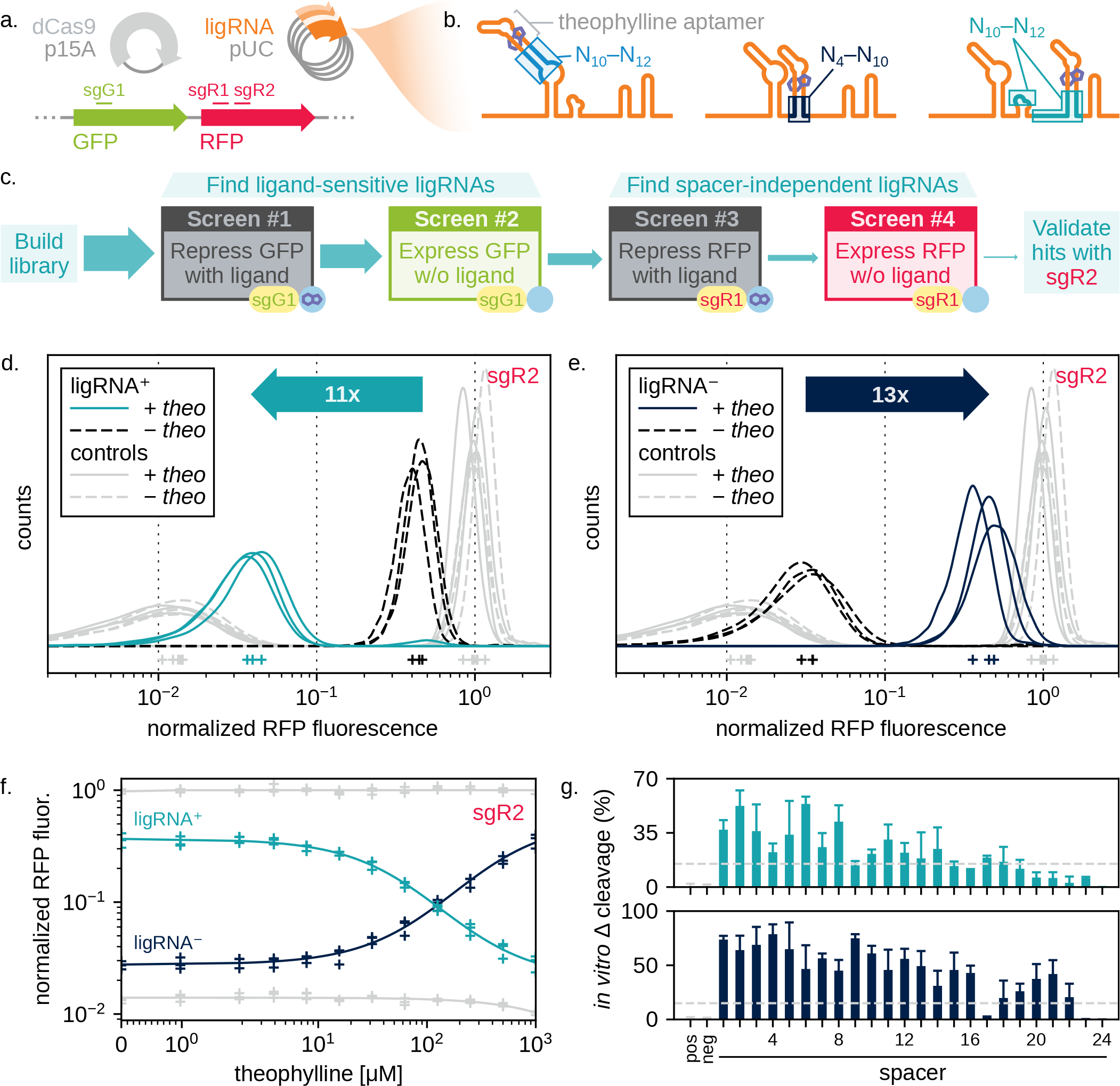
Identification of robust ligRNAs using CRISPRi-based gene repression in *E. coli.* (**a**) Components used in the CRISPRi assay. dCas9 and any ligRNAs were expressed from plasmids, while the fluorescent reporters (GFP and RFP) were chromosomally integrated. The DNA regions targeted by different spacers (sgG1, sgR1, sgR2) used to repress the fluorescent reporters are indicated. (**b**) Regions randomized in each ligRNA library. (**c**) Schematic of the screen used to isolate ligRNA^+^. sgG1, sgR1, and sgR2 refer to spacers targeting GFP and RFP, respectively (**Table S4**). (**d,e**) Single-cell RFP fluorescence distributions for ligRNA^+^ (teal, panel **d**) and ligRNA^−^ (navy, panel **e**) targeting RFP using the sgR2 spacer with (solid lines) and without (dashed lines) theophylline. Control distributions are in grey (positive control: optimized sgRNA scaffold[36]; negative control: G43C G44C[28]). The mode of each distribution is indicated with a plus sign. RFP fluorescence values for each cell are normalized by both GFP fluorescence for that cell and the modes of the unrepressed control populations (i.e. *apo* and *holo*) measured for that replicate. (**f**) Efficiency of CRISPRi repression with increasing theophylline concentrations for ligRNA^+^ (teal) and ligRNA^−^ (navy). Controls are in grey. The fluorescence axis is the same as in (**d**) and (**e**). The fits are to a two-state equilibrium model. (**g**) Change in the percentage of DNA cleaved *in vitro* in the presence and absence of theophylline for ligRNA^+^ and ligRNA^−^ in the context of 24 representative spacers. Each bar represents the mean of three or four measurements with a single spacer (except for spacer #24, where n=2, **Table S5**), and the error bars give the standard deviation. Pos and neg denote the positive and negative controls (**Table S4**), which are independent of theophylline.

**Figure 3:**
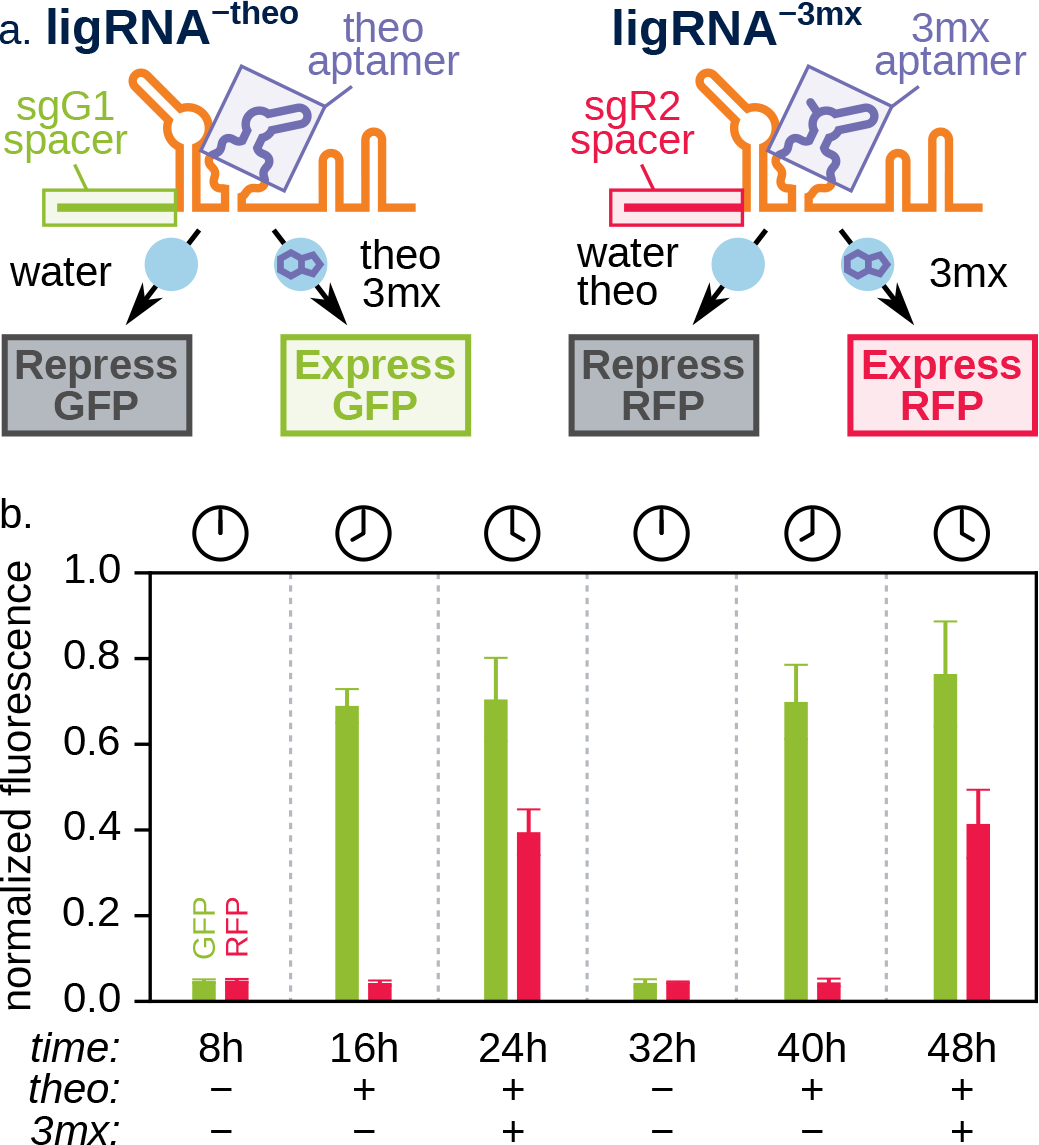
Multiplexed temporal control of two genes with two ligands. (**a**) Schematic illustrating the constructs and the expected consequences of adding theophylline (theo) and 3-methylxanthine (3mx) for each fluorescent reporter. (**b**) GFP and RFP fluorescence measured at indicated time points by flow cytometry. Presence of theo leads to GFP expression; addition of 3mx separately at a different time point also triggers RFP expression. Both effects are reversible (GFP and RFP are repressed when both ligands are absent) and expression can be triggered a second time with ligand addition. Bar heights and error bars represent the modes and standard deviations of the cell distributions, respectively. Unlike in Figure 2, fluorescence is normalized by side-scatter because both fluorescent channels are being manipulated.

## Contributions

KK conceived the idea for the project, KK and TK conceived the experimental approach, KK designed and performed the majority of the experiments and analyzed the data with advice and contributions from JEL (CRISPRi multiplexing with different ligands), KEW (ligRNA^−^ mechanism), BLO (FACS selections) and CMF (*in vitro* DNA cleavage experiments). CF carried out the mammalian cell line experiments. AHN and BMH carried out the yeast experiments. DFS, HES, JAD and TK provided advice, mentorship and resources. KK and TK wrote the manuscript with input from all authors.

## Acknowledgments

The authors would like to thank Anna Simon for discussing research that in part inspired the project, Kianna Zucker for generating constructs used in **Figure 2g**, and Kyle Barlow and Anum Glasgow for experimental help. The work was supported by National Institutes of Health (NIH) grant R21-EB021453 to TK and a Medical Research Award to TK by the W.M. Keck Foundation. JEL was supported by a Graduate Fellowship from the National Science Foundation (NSF GRFP). HES and TK are Chan Zuckerberg Biohub Investigators. CF is supported by a US National Institutes of Health K99/R00 Pathway to Independence Award (K99GM118909) from the National Institute of General Medical Sciences (NIGMS). BMH is supported by the Post 9/11 GI Bill. JAD is a Paul Allen Frontiers Group Distinguished Investigator and an HHMI Investigator.

## Methods

### Constructs

All experiments used Cas9 from S. *pyogenes* (called Cas9 throughout), with mutations D10A and H840A for CRISPRi experiments (dCas9). Sequences of relevant ligRNAs, aptamers, spacers and controls are listed in **Table S4**.

### *In vitro* DNA cleavage assay

<u>sgRNA *in vitro* transcription</u>: Linear, double-stranded template DNA was acquired either by ordering gBlocks^®^ Gene Fragments from IDT (experiments in Figure 1d,e) or by cloning the desired sequence into a pUC vector and digesting it with EcoRI and HindIII (experiments in Figure 2g). Each construct contained a T7 promoter and a spacer that began with at least 3 Gs (**Table S4**). DNA template (10–50 ng) was transcribed using the HiScribe™ T7 High Yield RNA Synthesis Kit (NEB E2040S) and unincorporated ribonucleotides were removed with Zymo RNA Clean & Concentrator™-25 spin columns (Zymo R1018).

<u>Target DNA</u>: Target DNA was prepared using inverse PCR to clone the appropriate sequence into a modified pCR2.1 vector approximately 2.1 kb downstream of its XmnI site. The vector was then digested with XmnI (NEB R0194S) as follows: mix 43.5 μL ≈500 ng/μL miniprepped pCR2.1 DNA, 5.0 μL 10X CutSmart buffer, and 1.5 μL 20 U/μL XmnI; incubate at 37°C until no uncleaved plasmid is detectable on a 1% agarose gel (usually 30-60 min); dilute to 30 nM in 10 mM Tris-Cl, pH 8.5; store at -20°C.

<u>Cas9 reaction</u>: We adapted the following protocol from ref. [28]: mix 5.0 μL water or 30 mM theophylline (in water) and 1.5 μL 1.5 μM sgRNA (in water); incubate at 95°C for 3 min, then at 4°C for 1 min; prepare Cas9 master mix for 40 reactions: 241.0 μL water, 66.0 μL 10x Cas9 buffer (NEB B0386A), and 1.0 μL 20 μM Cas9 (NEB M0386T); add 7.0 μL Cas9 master mix; incubate at room temperature for 10 min; add 1.5 μL 30 nM target DNA; pipet to mix; incubate at 37°C for 1 h; prepare quenching master mix: 4.68 μL 20 mg/mL RNase A (Sigma R6148), 4.68 μL 20 mg/mL Proteinase K (Denville CB3210-5), and 146.64 μL 6x Orange G loading dye via master mix; add 3 μL quenching master mix; incubate at 37°C for 20 min, then at 55°C for 20 min; run the entire reaction (18 μL) on a 1% agarose/TAE/GelRed gel at 4.5 V/cm for 70 min.

<u>Gel quantification</u>: Band intensities were quantified using Fiji (1.51r)[37]. The background was subtracted from each image using a 50 pixels rolling ball radius. The fraction of DNA cleaved in each lane (f) was calculated as follows (pixels_2kb_ and pixels_4kb_ are the intensities of the cleaved and uncleaved bands, respectively):

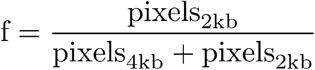

The change in cleavage due to ligand (Δf) was calculated as follows (f_apo_ and f_holo_ are the fractions of DNA cleaved in the reactions without and with theophylline, respectively):

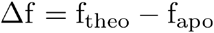

### CRISPR-Cas9-based repression (CRISPRi) assay in *E. coli*

<u>Strain</u>: The strain used for all CRISPRi experiments was *E. coli* MG1655 with dCas9 (containing the D10A and H840A mutations) and ChlorR on a p15A plasmid (pgRNA-bacteria, Addgene 44251), sgRNA and AmpR on a pUC plasmid (pdCas9-bacteria, Addgene 44249), and sfGFP[38], mRFP[39], and KanR chromosomally integrated at the *nsfA* locus. This strain was originally described in ref.[2].

<u>Flow cytometry</u>: Overnight cultures of the CRISPRi strain above were inoculated from freshly picked colonies in 1 mL Lysogeny Broth (LB) medium with 100 μg/mL carbenicillin (100 mg/mL stock in 50% EtOH) and 35 μg/mL chloramphenicol (35 mg/mL stock in EtOH). The next morning, fresh cultures were inoculated in 15 mL culture tubes or 24-well blocks by transferring 4 μL of overnight culture into 1 mL EZ Rich Defined Medium (Teknova M2105) with 0.1% glucose, 1 μg/mL anhydrotetracycline, 100 μg/mL carbenicillin, 35 μg/mL chloramphenicol, with or without 1 mM theophylline (added from a 30mM stock dissolved in water). These cultures were then grown for 8h at 37°C with shaking at 225 rpm before GFP (488 nm laser, 530/30 filter) and RFP (561 nm laser, 610/10 filter) fluorescence were measured using a BD LSRII flow cytometer. Approximately 10,000 events were recorded for each measurement. Biological replicates were performed on different days using different colonies from the same transformation.

<u>Data analysis</u>: Cell distributions were obtained by computing a Gaussian kernel density estimation (KDE) over the base-10 logarithms of the measured fluorescence values. The mode was considered to be the center of each distribution (e.g. for determining fold changes) and was obtained through the Broyden-Fletcher-Goldfarb-Shanno (BFGS) maximization of the KDE. Dose response curves were fit to the Hill equation (y is the normalized fluorescence, x in the theophylline concentration, EC50 is the inflection point, and y_min_ and y_max_ are the lower and upper asymptotes of the fit):

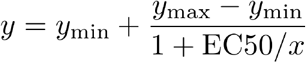

### Identification of functional ligRNAs in *E. coli* using FACS screens

<u>Library generation</u>: Randomized regions were inserted into the sgRNA using inverse polymerase chain reaction (PCR) with phosphate-modified and high-performance liquid chromatography (HPLC)-purified primers containing degenerate nucleotides.

<u>Electrotransformation</u>: Electrocompetent cells were prepared as follows: make “low-salt” Super Optimal Broth (SOB) medium: 20 g bacto-tryptone, 5 g bacto-yeast extract, 2 mL 5 M NaCl, 833.3 μL 3 M KCl, water to 1L, pH to 7.0 with NaOH, autoclave 30 min at 121°C; pick a fresh colony and grow overnight in 1 mL SOB; in the morning, inoculate 1 L SOB with the entire overnight culture; grow at 37°C with shaking at 225 rpm until OD=0.4 (≈4 h); place cells in an ice bath for 10 min; wash with 400 mL pre-chilled water, then 200 mL pre-chilled water, then 200 mL pre-chilled 10% glycerol; resuspend in a total volume of 6 mL pre-chilled 10% glycerol; make 100 μL aliquots; flash-freeze and store at - 80°C. Electrocompetent cells were transformed as follows: thaw competent cells on ice for 10 min; pipet once to mix cells with 2 μL ≈250 ng/μL library plasmid; shock at 1.8 kV with a 5 ms decay time; immediately add 1 mL pre-warmed SOB with catabolite repression (SOC) medium; recover at 37°C for 1 h; dilute into selective liquid media and grow at 37°C with shaking at 225 rpm overnight. After PCR and ligation, libraries were first transformed into electrocompetent Top10 cells, then mira-prepped[40], sequenced, and combined to achieve approximately equal representation of variants based on library size and DNA concentration, then transformed again into electrocompetent MG1655 cells already harboring the dCas9 plasmid.

<u>Cell sorting</u>: Cells were grown as for the CRISPRi assay, but when starting new cultures, care was taken to subculture at least 10x more cells than the size of the library (often 200 μL). Sorting was done using a BD FACSAria II cell sorter. Sorting was no slower than 1000 evt/s and no faster than 20,000 evt/s, with the slower speeds being more accurate and the faster speeds being necessary to sort large libraries. Gates were drawn based on the position of the control population if possible, and based on the most extreme library members otherwise. Typically the gates included between 1% and 5% of the population being sorted. All gates were drawn diagonally in GFP vs. RFP space. Sorted cells were collected in 1 mL SOC at room temperature and, after sorting, were diluted into selective media and grown at 37°C with shaking at 225 rpm overnight.

<u>Screening for ligRNA^+^</u>: Pool libraries 23-28 from **Table S2**. First screen: grow without ligand, gate for GFP expression, sort 10,000 evt/s for 3.5 h. Second screen: grow with ligand, gate for GFP repression, sort 1500 evt/s for 70 min. Third screen: grow without ligand, gate for GFP expression, sort 1700 evt/s for 10 min. Fourth screen: grow with ligand, gate for GFP repression, sort 1000 evt/s for 2 min. Fifth screen: grow without ligand, gate for GFP expression, sort 5000 evt. Plate cells and test 96 individual colonies using the CRISPRi assay. Miniprep and sequence the 20 selected designs with the largest response to theophylline. Only one unique sequence was identified, and it did not function with the sgR1 spacer (**Table S3**). Note that we did not change the spacer in between the second and third screens for this library. We then designed libraries 29-30 (**Table S2**) to keep the stem identified in the previous screen of libraries 23-28 and to randomize other regions of the sgRNA that might be participating in ligand-dependent base-pairing. First screen: grow with ligand, gate for GFP repression, sort 4000 evt/s for 2 h. Second screen: grow without ligand, gate for GFP expression, sort 1500 evt/s for 1 h. Change the spacer from sgG1 to sgR1 (**Table S4**) using inverse PCR. Third screen: grow with ligand, gate for RFP repression, sort 2000 evt/s for 10 min. Fourth screen: grow without ligand, gate for RFP expression, sort 10,000 evt. Plate cells and test 96 individual colonies using the CRISPRi assay. Miniprep and sequence the 15 selected designs with the largest response to theophylline, then test those designs with four different spacers (sgG1, sgR1, sgG2, sgR2). There was only 1 duplicate sequence, and 8 of the sequences had acquired unexpected mutations outside of the randomized region. The majority of these hits were tested with four different spacers (sgG1, sgR1, sgG2, and sgR2). ligRNA^+^ and ligRNA^−^ performed best (none of the hits performed well with the sgG2 spacer.) Details on all library hits and validation with different spacers can be found in **Table S3**.

<u>Screening for ligRNA^−^</u>: Pool libraries 7-22 from **Table S2**. First screen: grow without ligand, gate for GFP repression, sort 18,000 evt/s for 1 h. Second screen: grow with ligand, gate for GRP expression, sort 1000 evt/s for 10 min. Third screen: grow without ligand, gate for GFP repression, sort 1000 evt/s for 10 min. Fourth screen: grow with ligand, gate for GRP expression, sort 1000 evt/s for 10 min. Fifth screen: grow without ligand, gate for GFP repression, sort 1500 evt/s for 7 min. Plate cells and test 96 individual colonies using the CRISPRi assay described above. Miniprep and sequence the 20 selected designs with the largest fold response to theophylline. In this group there were only 9 unique sequences. ligRNA^−^ appeared 5 times and had the largest response to theophylline (**Table S3**). Note ligRNA^−^ is functional with other spacers (Figure 2g) despite the fact that we did not change the spacer in between the second and third screens for this library.

### Test of different spacers

The spacers for this assay were chosen by a script that generated uniformly random sequences, scored them using a previously published machine learning approach for designing functional sgRNAs[41], and kept only those that scored higher than 0.5 (the median). This approach was designed to produce spacers that were as unbiased as possible, while still being likely to function as expected in the positive (optimized sgRNA scaffold[36]) and negative (G43C G44C[28]) controls shown in **Table S4**. The average cleavage for the positive controls was 93%, and the lowest cleavage for any of the positive controls was 78% (**Table S5**).

### RNA secondary structure predictions

Secondary structure predictions were performed using the RNAfold program from the ViennaRNA package (version 2.4.3). The structures reported here are minimum free energy (MFE) predictions, although centroid and maximum expected accuracy (MEA) structures from partition function calculations were nearly identical in every case. The *holo* state was simulated using soft constraints: a -9.21 kcal/mol bonus was granted for forming the base pair flanking the aptamer. This bonus corresponds to the 320 nM affinity of the theophylline aptamer for its ligand[30].

The command-lines used for the *apo* and *holo* states, respectively, are given below:

$ RNAfold --partfunc --MEA
$ RNAfold --partfunc --MEA --motif \

> “GAUACCAGCCGAAAGGCCCUUGGCAGC,(…((((((….)))…)))…),-9.212741321099747”

Free energy predictions for **Figure S2** were performed using the RNAduplex program from the ViennaRNA package (version 2.4.3):

$ RNAduplex

### CRISPRi assay in *S. cerevisiae*

<u>Yeast Strains</u>. All yeast strains were constructed using the MoClo golden gate cloning framework and the Yeast Toolkit from[42]. The background strain was WCD230 (derived from BY4741 (MATa his3Δ1 leu2Δ0 met15Δ0 ura3Δ0) with a larger fraction of the his3 gene removed[43]. All yeast strains contained the dCas9-Mxi1 inhibitor, fluorescent reporters, and ligRNA or control sgRNA construct, respectively (**Figure S10a**). The inhibitor cassette was integrated into the Ura3 locus and contained pGal1-dCas9-Mxi1 and pRnr2-GEM constructs that allowed for estradiol-inducible expression of dCas9-Mxi1[44]. The Mxi1 inhibition domain was derived from Addgene catalog number 46921[3] with synonymous mutations made to the E68 and T69 codons to render the construct compatible with golden gate cloning. The fluorescent reporter cassette was integrated into the His3 locus and contained pCcw12-sfGFP and pTdh3-mRFP. The guide cassette was integrated into the Leu2 locus and contained (tRNAPhe)-HDV Ribozyme-sgRNA. The HDV Ribozyme cleaved the ligRNA or control sgRNA from the rest of the transcript to prevent unwanted interactions[45]. The sgRNA was either positive or negative control sgRNA, ligRNA^+^, ligRNA^+^_2_, or ligRNA^−^.

<u>Experimental protocol</u>: Strains were plated from freezer stocks on SD-HIS (Yeast Nitrogen Base (YNB), Complete Synthetic Media (CSM) lacking histidine, 2% glucose) because the fluorescent reporter cassette integrated into the His3 locus caused a slight growth defect that was also present for an empty vector integrated in the same locus. After incubating at 30°C, single colonies were grown overnight in 0.5ml YPD (yeast extract, peptone, 2% glucose) in 96 well plates with 2ml/well maximum capacity, shaking at 900 RPM at 30C in an Infors HT Multitron Pro shaker. Saturated cultures were diluted 1:100 into 1ml SDC (YNB, CSM, 2% glucose) in a new 2ml 96 well plate and placed at 30°C shaking at 900RPM for 2 hours. Cells were then diluted 1:4 into 400ul SDC with estradiol and either theophylline from a 30mM stock or water in a new 2ml 96 well plate. The final concentration of estradiol was 125nM and the final concentration of theophylline when present was 2.5mM. After 8 hours shaking at 900RPM and 30°C, cells were diluted in 1XTE buffer and analyzed on a flow cytometer (BD LSR II).

### Gene editing assay in mammalian cells

HEK293T cells, and derived cell lines, were grown in Dulbecco’s Modified Eagle Medium (DMEM; Corning Cellgro, #10-013-CV) supplemented with 10% fetal bovine serum (FBS; Seradigm #1500-500), and 100 Units/ml penicillin and 100 μg/ml streptomycin (Pen-Strep; Life Technologies Gibco, #15140-122) at 37°C with 5% CO_2_.

HEK293T-based genome editing reporter cells, referred to as HEK-RT1, were established in a two-step procedure. In the first step, puromycin resistant monoclonal HEK-RT3-4 reporter cells were generated as previously described (Hui Liu *et al.*, in revision). In brief, HEK293T human embryonic kidney cells were transduced at low-copy with the amphotropic pseudotyped retrovirus RT3GEPIR-sh.Ren.713[46], comprising an all-in-one Tet-On system enabling doxycycline-controlled EGFP expression. After puromycin (2.0 μg/ml) selection of transduced HEK239Ts, 36 clones were isolated and individually assessed for i) growth characteristics, ii) homogeneous morphology, iii) sharp fluorescence peaks of doxycycline (1 μg/ml) inducible EGFP expression, iv) relatively low fluorescence intensity to favor clones with single-copy reporter integration, and v) high transfectability. HEK-RT3-4 cells are derived from the clone that performed best in these tests. In the second step, HEK-RT1 cells were derived by transient transfection of HEK-RT3-4 cells with vectors encoding Cas9 and sgRNAs targeting puromycin, followed by identification of monoclonal reporter cell lines that are puromycin sensitive.

A lentiviral vector, referred to as pCF204, expressing a U6 driven sgRNA and an EFS driven Cas9-P2A-Puro cassette was based on the lenti-CRISPR-V2 plasmid[47], by replacing the sgRNA with an enhanced *Streptococcus pyogenes* Cas9 sgRNA scaffold[5]. All sgRNAs (sgRen71: TAGGAATTATAATGCTTATC, sgGFP1: CCTCGAACTTCACCTCGGCG, sgGFP9: CCGGCAAGCTGCCCGTGCCC) were designed with a G preceding the 20-nt guide for better expression, and cloned into the lentiviral vector using the BsmBI restriction sites. The lentiviral vectors expressing ligRNA^+^, ligRNA_2_^+^ and ligRNA^−^ (referred to as pCF441, pCF442 and pCF443, respectively) were all based on pCF204, by replacing the SpyCas9 sgRNA scaffold with the respective ligRNAs using custom oligonucleotides (IDT), gBlocks (IDT), standard cloning methods, and Gibson assembly techniques. Lentiviral particles were produced using HEK293T packaging cells; viral supernatants were filtered (0.45μm) and added to target cells. Transduced HEK-RT1 target cells were selected on puromycin (1.0μg/ml).

GFP expression in HEK-RT1 reporter cells was induced using doxycycline (1μg/ml; Sigma-Aldrich). Percentages of GFP-positive cells were assessed by flow cytometry (Attune NxT, Thermo Fisher Scientific), routinely acquiring 10,000-30,000 events per sample. Theophylline (Sigma-Aldrich, #T1633-50G) was used at the indicated concentrations, ranging from 0.1mM to 10mM. Note, theophylline concentrations of 5mM and 10mM resulted in considerable cellular toxicity in the HEK293T-based reporter cell line.

### Abbreviations

Cas9: CRISPR-associated protein 9
CRISPR: Clustered regularly interspaced short palindromic repeats
CRISPRi: CRISPR-inhibition
FACS: Fluorescence activated cell sorting
mRFP: Monomeric red fluorescent protein
PAM: Protospacer adjacent motif
sfGFP: Super-folder green fluorescent protein
sgRNA: single guide RNA

